# Effects of aqueous antioxidants extracts from ginger, garlic and onion on 2-thiobarbituric Acid Reactive Substances and total volatile basic nitrogen content in chevon (goat meat) and pork during frozen storage

**DOI:** 10.1101/2021.08.13.456289

**Authors:** Lesten Eliez Chisomo Chatepa, Kingsley George Masamba, Jonathan Tanganyika

**Affiliations:** Basic Science Department, Faculty of Agriculture, Lilongwe University of Agriculture and Natural Resources, Bunda Campus, P.O. Box 219, Lilongwe, Malawi; Food Science and Technology Department, Faculty of Food and Human Sciences, Lilongwe University of Agriculture and Natural Resources, Bunda Campus, P.O. Box 219, Lilongwe, Malawi; Animal Science Department, Faculty of Agriculture, Lilongwe University of Agriculture and Natural Resources, Bunda Campus, P.O. Box 219, Lilongwe, Malawi

**Keywords:** 2-thiobarbituric acid reactive substances (2-TBARS), total volatile basic nitrogen (TVB-N), antioxidants, pork, goat meat (chevon), ginger, garlic, onion

## Abstract

The study investigated the effect of 10% aqueous extracts of ginger (*Zingiber officinale* L.), garlic (*Allium sativum* L.) and onion (*Allium cepa* L.) on the quality of frozen chevon and pork as reflected by changes in values of 2-thiobartituric Acid Reactive Substances (2-TBARS), pH and total volatile basic nitrogen (TVB-N) over a 14 day storage under frozen conditions. Fresh samples each weighing 2 kg from chevon and pork were bought from the local slaughter houses in two locations 24hrs after slaughtering while ginger roots, garlic and onion bulbs were purchased from Mitundu local market in Lilongwe district, Malawi. The pH, total 2-TBARS and TVB-N of the thigh chevon and pork were measured from frozen storage at −20 °C after 14 days. The 10% aqueous extracts resulted in low pH values of 5.63, 5.79 and 5.67 at 14 d for chevon mixed with ginger, garlic and onion respectively. However, treated pork had higher pH content compared to treated chevon at 14 d. At day14 of frozen storage, the 2-TBARS expressed as mg MDA/kg meat, for chevon mixed with ginger, garlic and onion aqueous extracts were 2.62±0.01, 0 and 4.71±0.03 which was lower compared to the value of 5.93±0.01 for raw chevon. The TBARS values of chevon and pork mixed with ginger, garlic and onion and control chevon decreased from day 0 to 7 which eventually increased on 14 d. On 14 d, pork mixed with garlic extracts had lower TBARS value of 2.13±0.01 compared to 3.50±0.20, 2.26±0.01 and 3.92±0.01 for pork mixed with extracts of onion, ginger and control pork sample respectively. TVB-N, in mg/100g, was highest in control raw chevon and pork registering 95.70±0.32 and 84.00±0.40 at 14 d. Low values of TVB-N of 7.24±0.23, 12.37±0.23 and 16.61v±0.50 were registered in chevon mixed with aqueous extracts of ginger, garlic and onion compared to the values of 14.23±0.62, 22.87±0.47 and 18.86±0.14 for pork mixed with aqueous extracts of ginger, garlic and onion. The results of the study have revealed that natural aqueous antioxidant extracts of ginger, garlic and onion have antioxidative effect on lipid peroxidation in frozen stored fresh chevon and pork signifying that the use of these extracts can maintain quality.

## Introduction

Global financial value of the meat sector is projected to be at USD 1142.9 billion by 2023 from an estimated value of USD 945.7 billion in 2018 [1]. In 2020, global meat consumption was estimated at 360 million tons yearly with an increase of 58% in over the last decades. It is reported that population growth has caused 54% and the other 4% fraction comes from consumption per capita and increased consumers’ income [2]. The upsurge in the global meat production and consumption is defined by the high nutritive value which is significant for good human health [3].

Meat and meat based foods have high nutrient contents like vitamins, proteins, fats and minerals. Meat has high amount of n-polyunsaturated fatty acids and linoleic acid which are significant to human health [3]. Polyunsaturated fatty acids (PUFA) are important nutrients in human diet because they limit the initiation of cardiovascular diseases, hypertension and arthritis [4]. Lipids in meat and meat products enhances flavor, aroma, juiciness and tenderness [5]. The nutrients from meat are lost during ageing and shelf life and the preservation methods define the available nutrients for human consumption. The post-mortem ageing of meat results in flavor formation and protein break-down enhancing meat tenderness [6]. Traditionally, the carcass is hung on the rack to age without controlling the atmospheric temperature, airflow and humidity resulting in the loss of valuable nutrients [7]. The most commonly available preservation methods include freezing, drying, salting, fermentation and roasting [8, 9].

Thermal processing of meat improves shelf life, food safety, flavor and taste. However, application of heat causes adverse physical and chemical changes affecting nutritional value, flavor and the safety of the food [10]. Triacylglycerols in fats and oils from meat and meat products undergo oxidation reaction during frying, boiling and roasting in a process called lipid peroxidation. Lipid peroxidation of fats and oils produces primary oxidation products like hydroperoxides, epoxides, epidioxides and hydroxides [11, 12]. The primary oxidation products, further decompose into large amount of secondary and tertiary oxidation products like carbonyl compounds which includes aldehydes [13, 14]. The hydrolysis of hydroperoxide produces malondialdehydes (MDA), a secondary oxidation product whose concentration indicates the degree of lipid oxidation [15]. MDA causes off-flavor, changes in food taste [16, 17] and initiates the development of cardiovascular diseases (atherosclerosis), cancer and is a mutagen for it reacts with DNA [18, 19].

The quality of meat and meat products is compromised when frozen for a long period of time like losing color, undergoing lipid peroxidation and protein denaturation [20]. Oxidation of lipid and protein during freezing is associated with changes in flavor, texture and color which defines meat freshness and consumer acceptability [21]. Various authors have reported quality deterioration from frozen meat and meat products due to lipid peroxidation [22, 23]. Alterations in color, flavor and accumulations of carcinogenic primary oxidation products like hydroperoxides, radicals, aldehydes and epoxides has been reported during freezing of meat and meat products [24].

In frozen muscles of tissues of fish and meat, bacteria and enzyme action results in the production of volatile bases like ammonia, trimethylamine, dimethylamine and other volatile acid[25]. Proteins are broken down into ammonia and trimethylamine is a reduction product of trimethylamine oxide during meat spoilage under frozen shelf life [25]. Total volatile basic nitrogen (TVBN) has been used as an index for measuring meat freshness as ammonia, trimethylamine and dimethylamine content measurement [26, 27].

Application of synthetic antioxidants, like butylated hydroxyanisole, butylated hyroxytoluene (BHT) and propyl gallate, has been used to limit the initiation of lipid peroxidation in meat and meat products [28, 29]. However, synthetic antioxidants have been linked to carcinogenicity and other health safety issues calling for the use of natural antioxidants to either scavenge peroxy radical chains or limit the formation of free radicals [30, 31]. Despite the fact that synthetic antioxidants are safe when used with reference to relevant regulations like being applied as low as 0.02% w/w [32], meat industries have intensified research in natural antioxidants to meet consumers’ perception of health safety [33]. Natural antioxidants are extracted from plants with high concentrations of phytochemicals like phenolic compounds, ascorbic acids and carotenoids [34]. The natural antioxidants have been extracted from leaves, roots, stems, seeds, fruits and barks of many plants like ginger, rosemary, garlic and oranges [35, 36, 37].

Despite the global increase in the consumption of meat and meat products and the production of MDA in frozen meat, limited studies have been conducted in a majority of developing countries including Malawi on the application of natural antioxidants in frozen meat to limit the formation of MDA. Therefore, this study evaluated the effect of ginger, garlic and onion aqueous extracts on the quality of raw goat (chevon) meat and pork stored at frozen temperature of −20°C for 14 days. In this respect, changes in quality of the frozen goat meat and pork was determined by measuring 2-thiobartituric Acid Reactive Substances (2-TBARS), pH and total volatile basic nitrogen (TVB-N) over a 14 day period.

## Materials and methods

Fresh (chevon) goat meat and pork were purchased from the local butcheries at Mitundu and Nsabwe local trading centres in Lilongwe district, central Malawi. Fresh ginger, garlic and onion bulbs were purchased from Mitundu local market in Lilongwe district.

### Extraction and preparation of natural aqueous antioxidants extracts

The aqueous extract was prepared by following the method described by Ranendra et al. [38] with minor modifications. Ginger, garlic and onion bulbs were ground in an electric food blender and 20 g of the ground materials were mixed with 100 ml of distilled water in a beaker for 24 hrs at room temperature. The crude aqueous extracts were filtered using a Whatmann filter paper and were evaporated to quarter volume at 50°C in the drying oven.

### Sample preparation and chemical analyses

100 g of steaks of chevon and pork were immersed in 100 ml of 10% (v/v) solution of the ginger, garlic and onion extracts for 24 hrs in beakers. After 24 hrs, the samples were transferred into clean beakers and were frozen at 4°C for 24 hrs. The chemical analyses were conducted on day 0, 1, 3, 7 and 14 respectively.

### pH determination

The pH of the samples was measured by using a Kent pH meter in 10% (w/v) of the sample solution. 10 g of the sample was aseptically homogenised in 100 ml of distilled water, decanted and pH was measured [37].

### Total Volatile Basic-Nitrogen (TVBN) determination

Total Volatile Basic-Nitrogen was determined by using distillation method following the procedure described by Pearson [16] (1976). 10 g of the sample was homogenised in 100 ml of distilled water in 250 ml quick fit flask, 1 g of magnesium oxide was added and the mixture was distilled for 15 minutes into 25 ml of 4% boric acid solution with 2 drops of mixed indicator. The distillate was titrated against 0.1M hydrochloric acid (HCl) [16] and calculated as mg TVBN /100g of sample as follow;

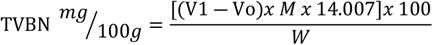

where V_1_ and V_o_ are HCl titre volumes of sample and blank respectively, M is the molar concentration of HCl.

### Thiobarbituric Acid Reactive Substances (TBARS) value determination

Thiobarbituric Acid Reactive Substances (TBARS) content was determined with reference to the distillation method of Tarladgis et al. [39] as described by Torres-Arreola et al [40] with minor modifications. 10g of the sample was minced into small pieces, transferred into a 250 ml quick fit flask and 47.5 ml and 2.5 ml of distilled water and 4 N HCl solution were respectively added. The mixture was swirled and distilled to collect 50 ml of the distillate. 5 ml of the distillate was pipetted into capped test tubes and 5 ml of 0.288% (w/v) solution of 2-thiobarbituric acid (TBA) in 50% (v/v) glacial acetic acid was added, the test tubes were capped and then boiled in a hot water bath for 35 minutes. Standard 1, 3, 3, 3 – tetramethoxypropane (TMP) samples of 0, 1 x10^-6^, 2 x10^-6^, 4×10^-6^, 8×10^-6^, 1.6 x 10^-6^ moles were prepared from a stock solution of 10^-6^ mole/litre by pipetting 0-6 ml into the test tubes. 5 ml of 2-TBA solution was added and the volume was made up to 10 ml with distilled water and were boiled for 35 minutes as the samples. The optical density was spectrophometrically measured at 532 nm using a UV-spectrophotometer. TBARS content as malondialdehyde (MDA), in mg/g, was calculated from the linear equation Y= 105742x as shown in Fig 1.

**Fig 1:**
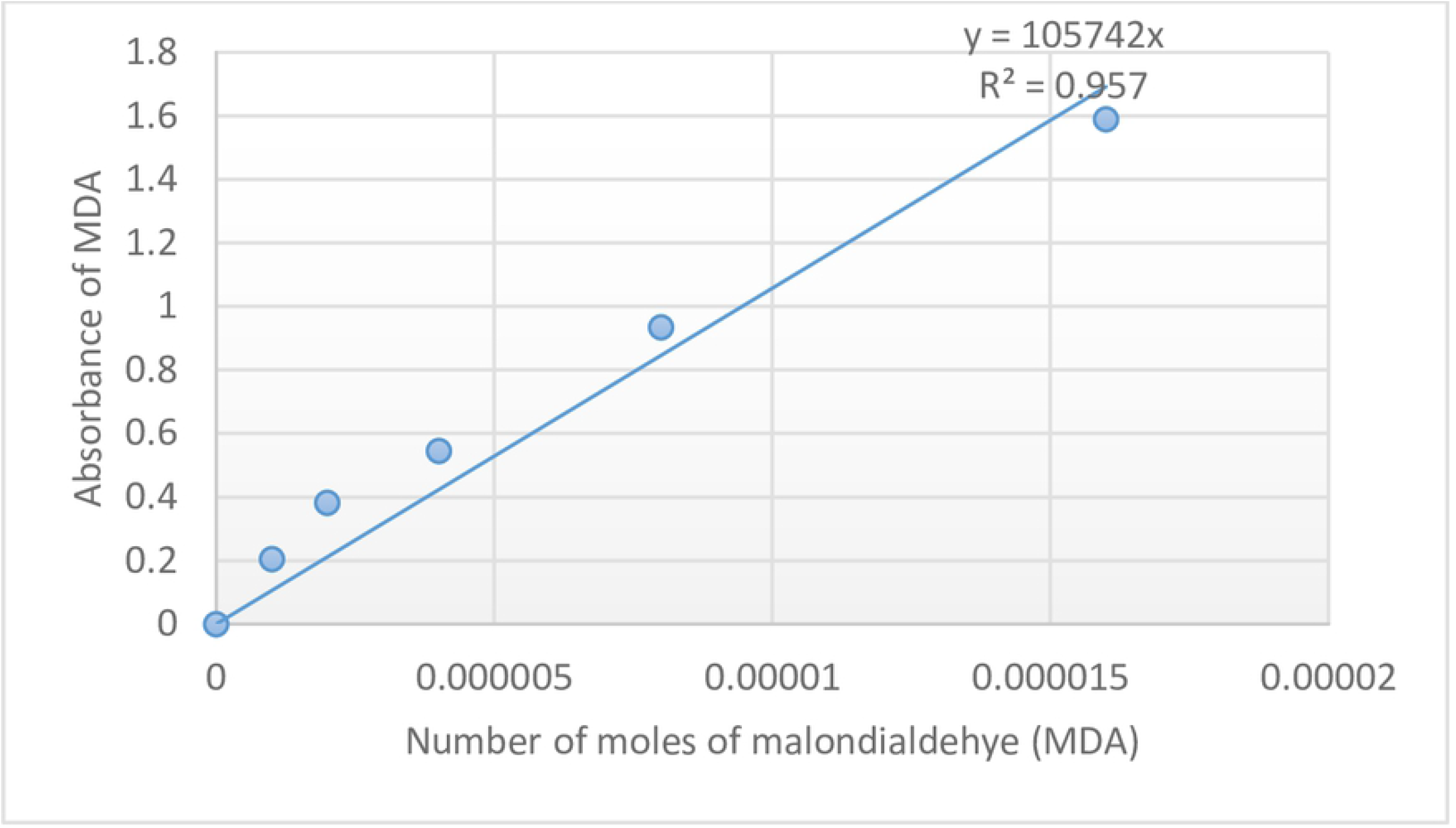
Standard curve of absorption of MDA against concentration

### Statistical Analysis

Laboratory chemical analyses were done in triplicate and the mean ± (SE) value of each chemical parameter was calculated using IBM SPSS version 20. The data was statistically analysed by using analysis of variance (ANOVA). T-test two-sample and unequal variances was used to compare mean values between treatments and meat type and significance was accepted at P≤ 0.05 level.

## Results and discussion

### pH changes of frozen chevon and pork with different treatments

Results on the pH changes of control and different treatment samples are presented in Fig 2. The initial pH values of control samples were 6.15 and 6.12 for chevon and pork respectively. The pH of the various treatment samples were 6.20, 5.48 and 6.40 and 6.76, 6.02 and 6.50 for raw/fresh chevon and pork mixed with 10% aqueous extracts of ginger, garlic and onion respectively. The pH of both control and treated samples increased after 3 days of frozen storage. However, the changes in pH slightly increased for both the controls and various treatments after 3 days of frozen storage which started decreasing after 7 days of storage. The lowest pH values of 5.63, 5.79 and 5.67 were observed in chevon compared to 6.67, 6.83 and 6.99 for pork mixed with 10% aqueous extracts of ginger, garlic and onion after 14 days of frozen storage respectively.

**Figure 2:**
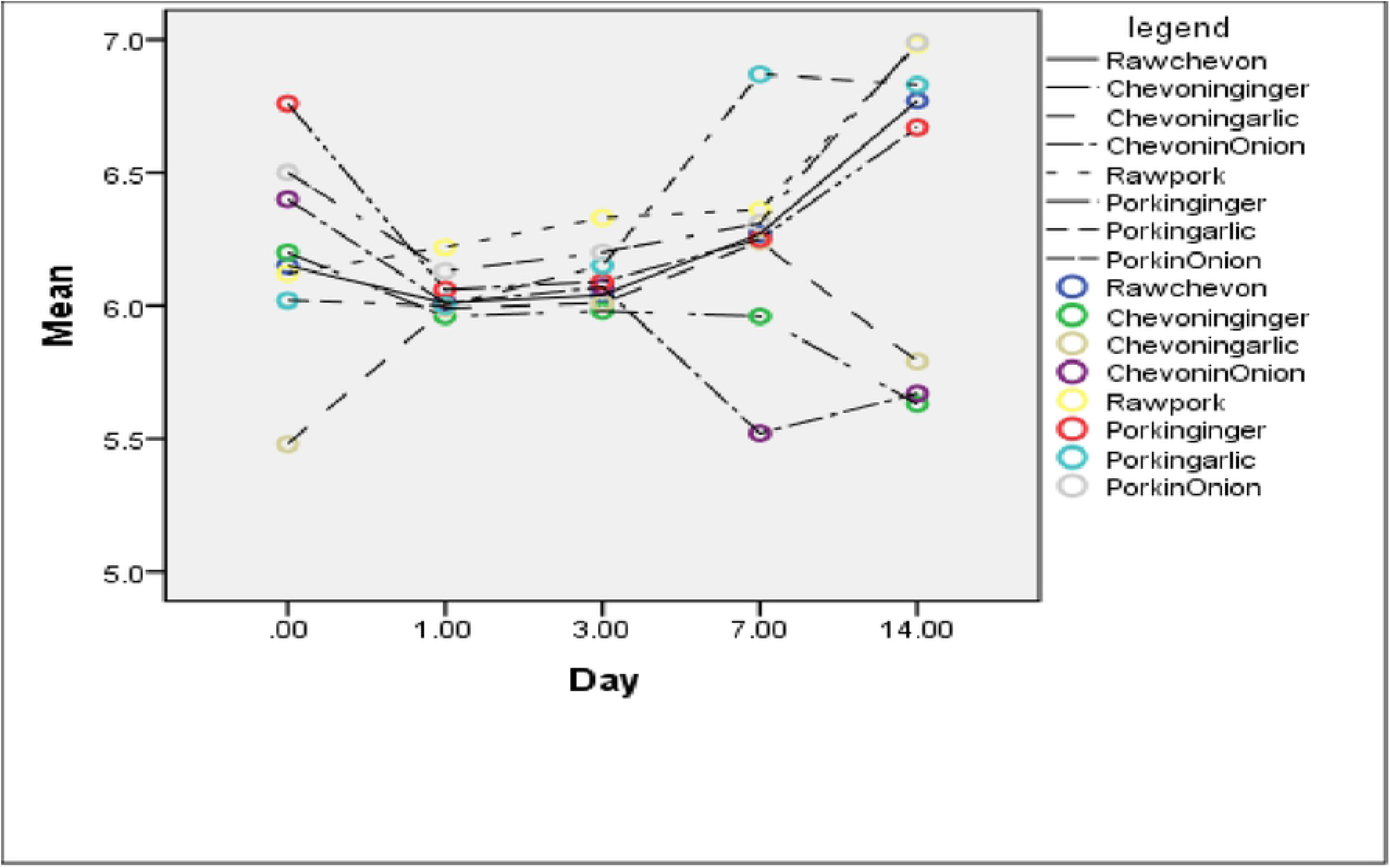
Changes in pH value of chevon and pork with different treatments during frozen storage at −4 °C Error bars: +/− 2 SE

The lowering/decreased of pH values has been attributed to the presence of lactic acid bacteria in meat samples [41]. Garlic and onions are reported to support the growth of lactic acid bacteria like *Lactobacillus* spp., *Weissella* spp. and *Leuconostoc* spp. [42]. The results from this study indicated that the lactic acid bacteria proliferated after day 1 of frozen storage which continuously kept the low pH of the samples up to day 14 of storage. The increase in the pH values after day 1 of storage could be attributed to protein degradation which results in amines production by micro-organisms in the meat [43]. It has been reported that the rise in pH of frozen stored meat is because of the production of volatile bases by endogenous or microbial enzymes. Alkaline ammonia like trimethylamine and ammonia are produced from amino acids after microbial decomposition [44]. In this study, the observed low pH values could be attributed to the antioxidants present in the natural herbal spices which suppressed the growth of basic nitrogen metabolizing microbes in the meat [45].

### 2-thiobarbituric reactive substances (2-TBARS) content in frozen chevon and pork

Results on 2-thiobarturic Reactive Substances as malondialdehyde (MDA) content (mg/kg) for chevon and pork samples are shown in Table 1. Meat undergo lipid oxidation when the unsaturated fat and protein are exposed to molecular oxygen besides the processing conditions [11, 12] producing secondary oxidation products like aldehydes resulting in off-flavours in meat and meat products [45]. In control chevon, TBARS content increased from 5.263 to 5.93 after 14 days of frozen storage. TBARS content in chevon treated with 10% aqueous extract of ginger, garlic and onion, decreased from 3.19±0.01, 4.01±0.01 and 4.47±0.00 to 2.03±0.01, 3.53±0.01 and 1.71±0.01 after 7 days of frozen storage but started increasing to day 14 of frozen storage. However, chevon mixed with 10% aqueous garlic extract had lower TBARS content of 1.45±0.01 followed by chevon treated with 10% aqueous onion extract (2.62±0.01) and chevon treated with 10% aqueous garlic (4.71±0.03) extracts at 14 d respectively. In pork, similar trend as that of chevon samples was observed and TBARS values of 10% aqueous extract of ginger, garlic and onion decreased to 1.78±0.01, 2.03±0.01 and 1.59±0.00 with increasing days of frozen storage up to day 7. Pork in 10% aqueous garlic extract had the lowest TBARS value of 2.13±0.01 followed by pork in 10% aqueous onion extracts (3.50±0.20 mg/kg) and 10% aqueous ginger extracts (2.26±0.01) respectively. However, the final TBARS contents after 14 days of frozen storage for pork treated with ginger, garlic and onion extracts were lower with the reported values of 2.26±0.01, 2.13±0.01 and 3.50±0.20 compared to 3.92±0.01 for control pork sample respectively.

**Table 1:**
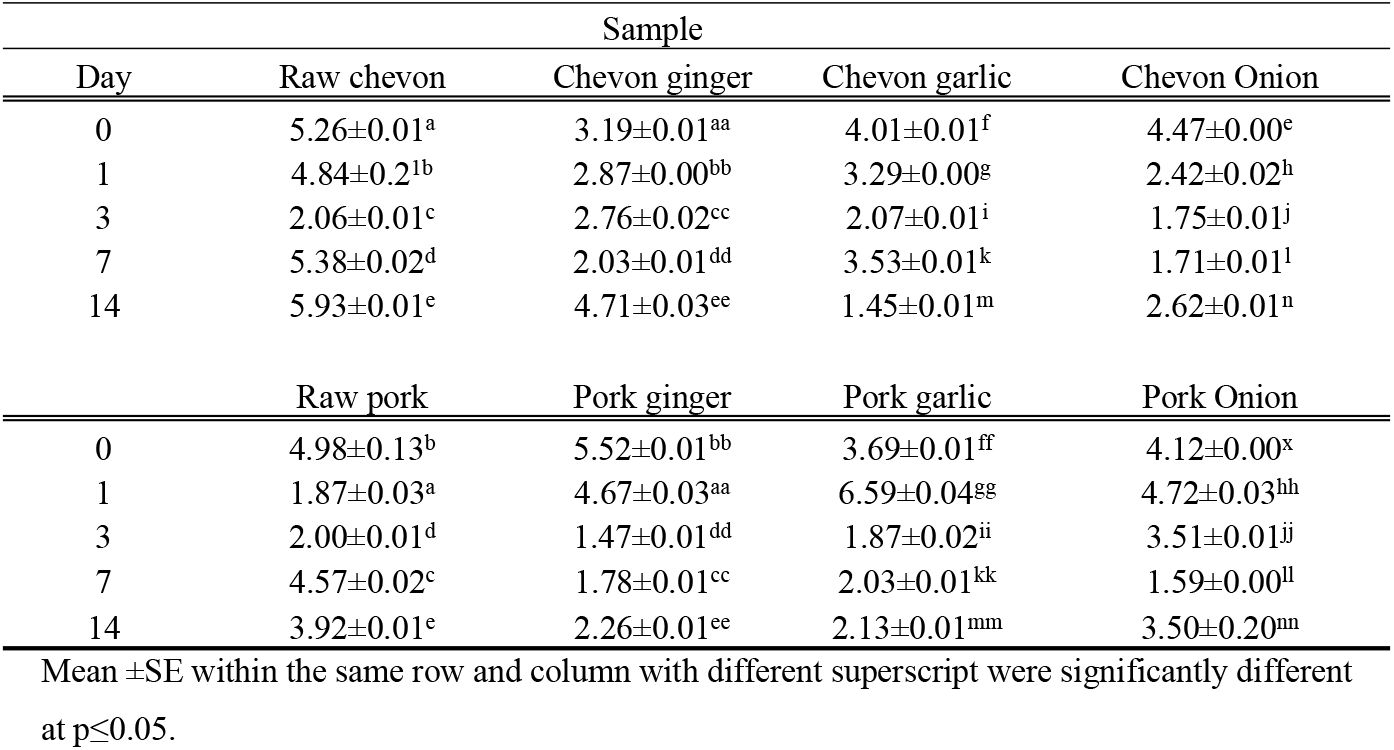
Mean composition of 2-thiobarbituric acid Reactive Substances (2-TBARS) [Mean ±SE] as MDA (mg/kg)

| Day | Sample |  |  |  |
| --- | --- | --- | --- | --- |
|  | Raw chevon | Chevon ginger | Chevon garlic | Chevon Onion |
| 0 | 5.26±0.01<sup>a</sup> | 3.19±0.01<sup>aa</sup> | 4.01±0.01<sup>f</sup> | 4.47±0.00<sup>e</sup> |
| 1 | 4.84±0.2<sup>1b</sup> | 2.87±0.00<sup>bb</sup> | 3.29±0.00<sup>g</sup> | 2.42±0.02<sup>h</sup> |
| 3 | 2.06±0.01<sup>c</sup> | 2.76±0.02<sup>cc</sup> | 2.07±0.01<sup>i</sup> | 1.75±0.01<sup>j</sup> |
| 7 | 5.38±0.02<sup>d</sup> | 2.03±0.01<sup>dd</sup> | 3.53±0.01<sup>k</sup> | 1.71±0.01<sup>l</sup> |
| 14 | 5.93±0.01<sup>e</sup> | 4.71±0.03<sup>ee</sup> | 1.45±0.01<sup>m</sup> | 2.62±0.01<sup>n</sup> |
| Day | Sample |  |  |  |
|  | Raw pork | Pork ginger | Pork garlic | Pork Onion |
| 0 | 4.98±0.13<sup>b</sup> | 5.52±0.01<sup>bb</sup> | 3.69±0.01<sup>ff</sup> | 4.12±0.00<sup>x</sup> |
| 1 | 1.87±0.03<sup>a</sup> | 4.67±0.03<sup>aa</sup> | 6.59±0.04<sup>gg</sup> | 4.72±0.03<sup>hh</sup> |
| 3 | 2.00±0.01<sup>d</sup> | 1.47±0.01<sup>dd</sup> | 1.87±0.02<sup>ii</sup> | 3.51±0.01<sup>jj</sup> |
| 7 | 4.57±0.02<sup>c</sup> | 1.78±0.01<sup>cc</sup> | 2.03±0.01<sup>kk</sup> | 1.59±0.00<sup>ll</sup> |
| 14 | 3.92±0.01<sup>e</sup> | 2.26±0.01<sup>ee</sup> | 2.13±0.01<sup>mm</sup> | 3.50±0.20<sup>nn</sup> |

Mean ± SE within the same row and column with different superscript were significantly different at $p \leq 0.05$ .

Cao et al., [45] reported low TBA value of 1.60 mg /kg for stewed pork samples which increased to 4.81 mg/kg after 12 days of refrigeration. However, low values of 0.43±0.01 and 0.40±0.01 mg MDA/kg meat for old female and young goat were reported in India [47]. Kim et al., [48] reported decreasing TBARS values of 0.083±0.002 and 0.074±0.001 mg/kg meat for pork marinated with 3 and 6% ginger extracts compared to 9.5±0.2 mg/kg meat for the control pork samples. Similarly, 3 and 6% onion juices had 0.084±0.001 and 0.080±0.001 TBARS content compared to 0.095±0.002 for control pork samples [48]. In pork patties mixed with fresh garlic, low TBARS values of 1.49 mg/kg have been reported and this value was lower compared to 1.92 mg/kg for control pork patties [35].

The findings from this study have shown that 2-TBARS values of 10% aqueous ginger, garlic and onion extracts mixed with chevon and pork samples were lower as compared to those of the control samples from day 0 to day 14 of frozen storage. This meant that antioxidants from ginger, garlic and onion were effective in reducing the extent of lipid oxidation in the frozen samples. Ginger, garlic and onion are reported to contain phenolic compounds, which are antioxidative [49] and could limit lipid oxidation during frozen storage of the meat samples [37]. The high TBARS values in control chevon of 5.93±0.01 compared to 3.92±0.01 for pork, indicates that animal species is one of the factors that affect lipid peroxidation in the carcass shelf life [50]. Lipid peroxidation is more susceptible in meat with high concentration of iron and myoglobin like raw beef and chevon compared to pork [51, 52].

### Total Volatile Basic Nitrogen (TVB-N) content of frozen chevon and pork

The composition of TVB-N values, in mg/100g, during frozen storage is presented in Table 2. Total volatile basic nitrogen (TVB-N) is a measure of the concentration of ammonia and primary, secondary and tertiary amines which defines the freshness of meat / muscles during storage [53]. TVB-N values of chevon during frozen storage increased up to 3 d but decreased up to 14 d. However, the TVB-N values of treatment samples was less than those of control samples from 7 d to 14 d. At 14 d, chevon mixed with 10% aqueous ginger extract had lower TVB-N value of 7.24±0.23 compared to 95.70±0.32, 12.37±0.23 and 13.77±0.23 for control and chevon mixed with 10% aqueous garlic and onion extract respectively.

**Table 2:**
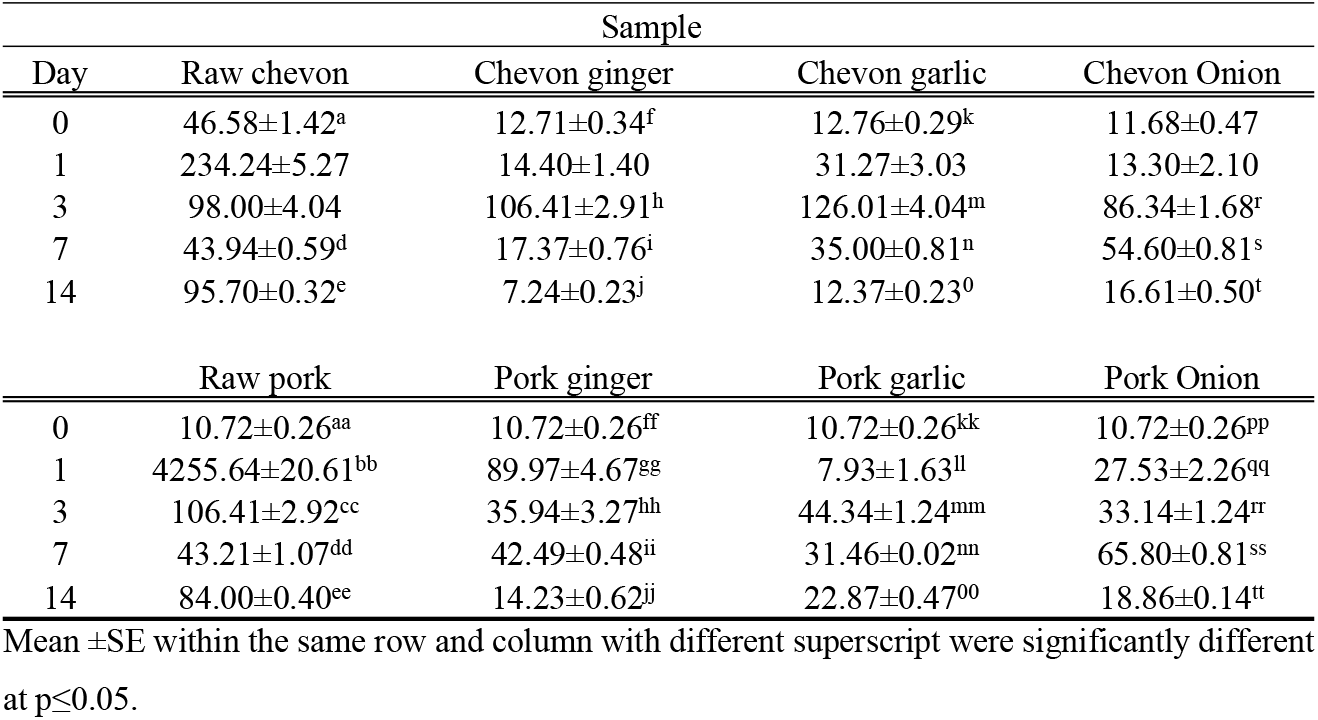
Total Volatile Basic Nitrogen (TVBN) [Mean ±SE] composition in mg/100 g

In pork, the TVB-N values of control and pork mixed with 10% aqueous ginger, garlic and onion extract were decreasing during the 14 days of frozen storage. At 14 d of frozen storage, pork mixed with 10% aqueous ginger extract had lower TVB-N value of 14.23±0.62 compared to 84.00±0.40 and 18.86±0.12 and 22.87±0.47 for pork mixed with 10% aqueous onion and ginger extracts respectively. The TVB-N value obtained in this study of 22.87±0.47 for pork mixed with 10% aqueous garlic extract was lower compared to 25, 22.5 and 15 mg/100 g meat for pork patties mixed 0.5% freeze-dried, freeze-dried fermented and freeze-dried aged garlic extract in a similar study conducted in Korea [54].

The increase in TVB-N values during storage of animal muscles is related to protein and amino acid breakdown by proteolytic gram-negative bacteria and enzymes [55]. It is reported that the presence of microbes like lactic acid bacteria, *Brochothrix thermosphacta, Enterobacteriaceae* and *Pseudomonas* increases bio-chemical reactions in meat during storage resulting in high concentration of TVB-N [56].

However, *Allium* species like garlic and onion have high concentration of phenolic compounds, sulphur and phenolic compounds which have anti-fungal and anti-microbial properties [57, 58, 59]. The hydroxyl groups in *Allium* spp reacts with either the sulfhydryl groups or some proteins of the bacteria limiting their proliferation in meat during storage [58, 59]. *Zingiber offinale* contains gingerol which has antimicrobial and antioxidant properties [60, 61] and therefore has the ability of limiting microbial growth in meat during frozen storage thereby reducing both TVB-N and TBARS values.

## Conclusion

The findings from the present study have shown that aqueous extracts of ginger, garlic and onion have antioxidative properties which inhibits lipid peroxidation in fresh chevon and pork during frozen storage. Application of 10% aqueous extracts resulted in low pH values compared to the control samples. Chevon and pork samples treated with 10% aqueous ginger, garlic and onion extracts had low TBARS and TVB-N values compared to the fresh control samples. Therefore aqueous extracts from ginger, garlic and onions could be used to maintain quality and subsequently prolong shelf life of frozen chevon and pork to limit the development of TBARS and TVB-N which initiate the development of cancer in human beings.

## Author contributions

**Conceptualization:** Lesten Eliez Chisomo Chatepa

**Data curation:** Lesten Eliez Chisomo Chatepa

**Formal analysis:** Lesten Eliez Chisomo Chatepa, Kingsley George Masamba

**Investigation:** Lesten Eliez Chisomo Chatepa, Kingsley George Masamba

**Writing - original draft:** Lesten Eliez Chisomo Chatepa, Kingsley George Masamba, Jonathan Tanganyika

**Writing - review & editing:** Lesten Eliez Chisomo Chatepa, Kingsley George Masamba, Jonathan Tanganyika

